# Visualization of Arabidopsis root system architecture in 3D by refraction-contrast X-ray micro-computed tomography

**DOI:** 10.1101/2021.05.04.442685

**Authors:** Tomofumi Kurogane, Daisuke Tamaoki, Sachiko Yano, Fumiaki Tanigaki, Toru Shimazu, Haruo Kasahara, Daisuke Yamauchi, Kentaro Uesugi, Masato Hoshino, Seiichiro Kamisaka, Yoshinobu Mineyuki, Ichirou Karahara

**Affiliations:** Graduate School of Science and Engineering for Education, University of Toyama, 3190 Gofuku, Toyama 930-8555, Japan; Faculty of Science, University of Toyama, 3190 Gofuku, Toyama 930-8555, Japan; Japan Aerospace Exploration Agency, 2-1-1 Sengen, Tsukuba 305-8505, Japan; Japan Space Forum, 3-2-1 Kandasurugadai, Tokyo 101-0062, Japan; Japan Manned Space Systems Corp., 1-1-26 Kawaguchi, Tsuchiura 300-0033, Japan; Graduate School of Life Science, University of Hyogo, 2167 Shosha, Himeji, Hyogo 671-2280, Japan; Japan Synchrotron Radiation Research Institute, 1-1-1 Kouto, Sayo, Sayo-gun, Hyogo 679-5198, Japan

**Author notes:** To whom correspondence should be addressed., Ichirou Karahara, Department of Biology, Faculty of Science, University of Toyama, 3190 Gofuku, Toyama, 930-8555, Japan.

**Keywords:** Arabidopsis, root system architecture, SPring-8, 3D observation, X-ray micro-CT, synchrotron radiation

## Abstract

Plant roots change their morphological traits in order to adapt themselves to different environmental conditions, resulting in alteration of the root system architecture. To understand this mechanism, it is essential to visualize morphology of the entire root system. To reveal effects of long-term alteration of gravity environment on root system development, we have performed an experiment in the International Space Station using Arabidopsis (*Arabidopsis thaliana* (L.) Heynh.) plants and obtained dried root systems grown in rockwool slabs (mineral wool substrate). X-ray computer tomography (CT) technique using an industrial X-ray scanner has been introduced for the purpose to visualize root system architecture of crop species grown in soil in 3D non-invasively. In the case of the present study, however, root system of Arabidopsis is composed of finer roots compared with typical crop plants and rockwool is also composed of fibers having similar dimension to that of the roots. A higher spatial resolution imaging method is required for distinguishing roots from rockwool. Therefore, in the present study, we tested refraction-contrast X-ray micro-CT using coherent X-ray optics available at the beamline BL20B2 of the synchrotron radiation facility SPring-8. Using this technique, both the primary and the secondary roots were successfully identified in the tomographic slices, clearly distinguished from the individual rockwool fibers and resulting in successful tracing of these roots from their basal regions. This newly-developed technique should contribute to elucidate the effect of microgravity on Arabidopsis root system architecture in space.

## Introduction

The plant root system, i.e., the belowground part of the plant body, provides a basis of supporting and anchoring the shoot system, i.e., the aboveground part of the plant body, as well as of growth by uptaking water and nutrients from soil. The plant root system adapts itself to surrounding soil environment changing its architecture [1]. This capability of the plant root system is called root system plasticity [2]. Understanding the mechanism of root plasticity is important for optimization of plant cultivation conditions under given environment. Greater root system development under mild drought stress is considered to contribute to their increased dry matter production [3]. Genotypes having increased total root length showed greater shoot dry matter production under mild water deficit condition [4].

The first step to understand mechanisms of root plasticity under different environmental stimuli is to precisely describe the morphology of the root system architecture. And the second one is to extract necessary information from it as much as possible. The methodology to analyze the root system architecture is still developing. A widely-used conventional method is to use rhizotron [5], which is to visualize the root system architecture *in situ* in 2D. On the other hand, modern technologies, such as, X-ray CT [6–11] [12] and magnetic resonance imaging (MRI) [13], are developed to visualize the root system architecture. For examples, root length data of the root system of 29-d old wheat plant (*Triticum aestivum* L.) obtained by X-ray CT was compared with the data obtained using flatbed scanner, demonstrating high correlation between them. Metzner et al. (2015) examined the root system architecture of 23-d old bean plant (*Phaseolus vulgaris* L.) using both X-ray CT and MRI and compared the data. In most of the previous studies, an industrial CT scanner or a scanning system employing microfocus X-ray source is used, which is suitable for routine analyses for the size of root systems of crop plants.

We have performed an experiment in the International Space Station ‘Kibo’ module called “Space Seed” to understand effects of Earth’s gravity on the entire life cycle of Arabidopsis plants[14] and obtained dried root systems grown in rockwool slabs. For this case, we have to deal with Arabidopsis root system which is composed of finer roots than those of crop plant species as mentioned above as well as to distinguish the roots from rockwool fibers having similar dimension to that of the roots. Therefore, we focused on refraction-contrast X-ray micro-CT technique, which is available at the beamline of the synchrotron radiation facility SPring-8, because high brightness of nearly-parallel X-ray beams enables imaging of samples at the micron scale [15]. In the present study, we tested two different Hutches available at this beamline and aimed to optimize the observation condition for the visualization of dried Arabidopsis root system developed in the rockwool slab.

## Methods and materials

### Plant materials and growth conditions

The present experiment was performed during the preparation of the Space Seed experiment. Growth conditions are basically the same as described previously [14], while the plant materials and the employed instrument were as follows. Twenty-four seeds of Arabidopsis (*Arabidopsis thaliana* (L.) Heynh.) Landsberg *erecta* (Ler) were sterilized and sown on a rockwool slab (W × D × H = 50 × 42 × 10 mm) (Nichias Corp, Tokyo, Japan) using gum arabic and germinated in a polycarbonate growth chamber having the outer dimensions of W × D × H = 60 × 50 × 60 mm and the dimensions of its inner void space was W × D × H = 56 × 46 × 48 mm (Fig. 1). The rockwool slab was covered with a transparent plastic plate and growth chambers were installed in the prototype of the Plant Experimental Units, which was designed for experiments in the International Space Station [14]. Plants were illuminated laterally with LED matrix [16] and light intensity was 29 μmol m^-2^ s^-1^ when measured at the bottom center of the growth chamber. Plants used for the following experiment were grown for 46 days and the rockwool slabs were dried and the shoot system of the plants was removed before the observation. Typical plants during its development are shown in Fig. 1.

**Fig. 1.**
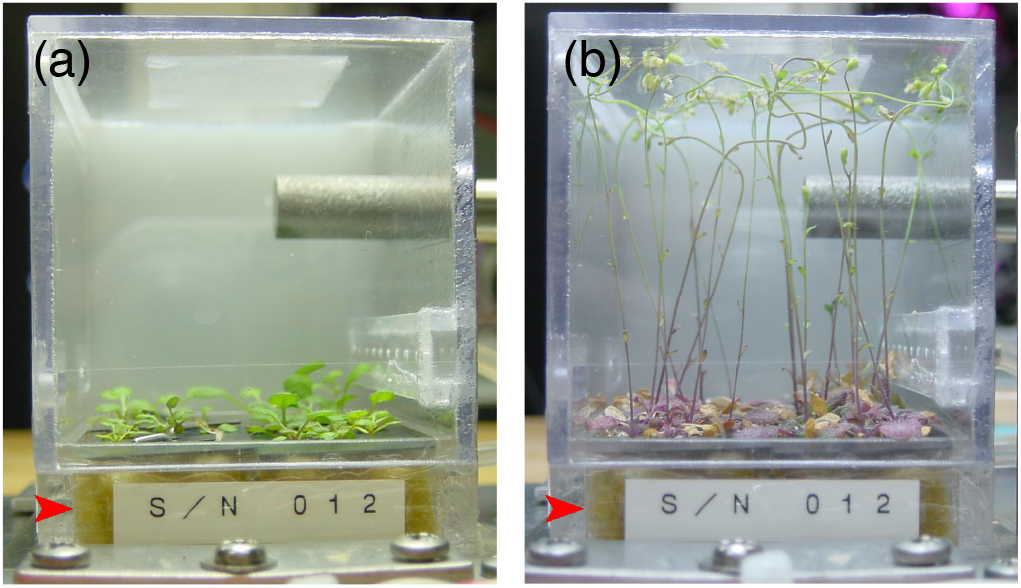
Pictures of typical Arabidopsis plants grown on a rockwool slab in a polycarbonate growth chamber. (a) Plants on Day 12. Rosette leaves are growing. Surface of the rockwool slab is seen underneath the plants. (b) Plants on Day 32. Flowers are forming. Internal length of the front side of the chamber is 46 mm. Red arrowheads indicate the rockwool slabs.

### Refraction contrast X-ray micro-CT

Refraction contrast X-ray micro-CT was performed at the experimental hutches (Hutch 1 and 3), where different spatial resolutions are available, of the beamline BL20B2 of the SPring-8 synchrotron radiation facility at Japan Synchrotron Radiation Research Institute, according basically to the method described by Karahara et al. [17]. Its experimental setup is shown in Fig. 2 and a brief of the method is as follows. The Hutch 1 and 3 are located 42 m and 206 m, respectively, from the bending magnet X-ray source. The X-ray energy was adjusted to 25 keV. The images consecutively projected on the fluorescent screen were recorded by a CMOS camera (ORCA-Flash 4.0; Hamamatsu Photonics KK, Hamamatsu, Japan) (Fig. 2a). The image sizes obtained at the Hutch 1 and Hutch 3 were 2048 × 2048 pixels (ca. 5 × 5 mm) and 2048 × 556 pixels, (ca. 50 × 15 mm), respectively. A series of 900 and 3000 projections were recorded over 180 degree for Hutch 1 and 3 observation, respectively. Because thickness of the rockwool slab was 10 mm, observation was separately done for the upper and lower halves for Hutch 1 observation. The spatial (pixel) resolution of the 3-D structure was estimated to be 25.5 μm pixel^-1^ for the Hutch 3 data, and 2.76 μm pixel^-1^ for the Hutch 1 data, respectively. The convolution back projection method was used for tomographic reconstruction [18] (Fig. 3b). To determining optimal filter used for reconstruction, Chesler, Ram-Lak, and Shepp-Logan filter, which are provided by the software package SP-μCT (http://www-bl20.spring8.or.jp/xct/), were tested and the results were compared from the aspect of background noises and clarity of root boundaries. Tomographic slices were obtained and 3-D models (isosurface, wireframe) were drawn using the IMOD software package [19], as previously described [20]. Volume models were drawn using the software UCSF Chimera (https://www.cgl.ucsf.edu/chimera/).

**Fig. 2.**
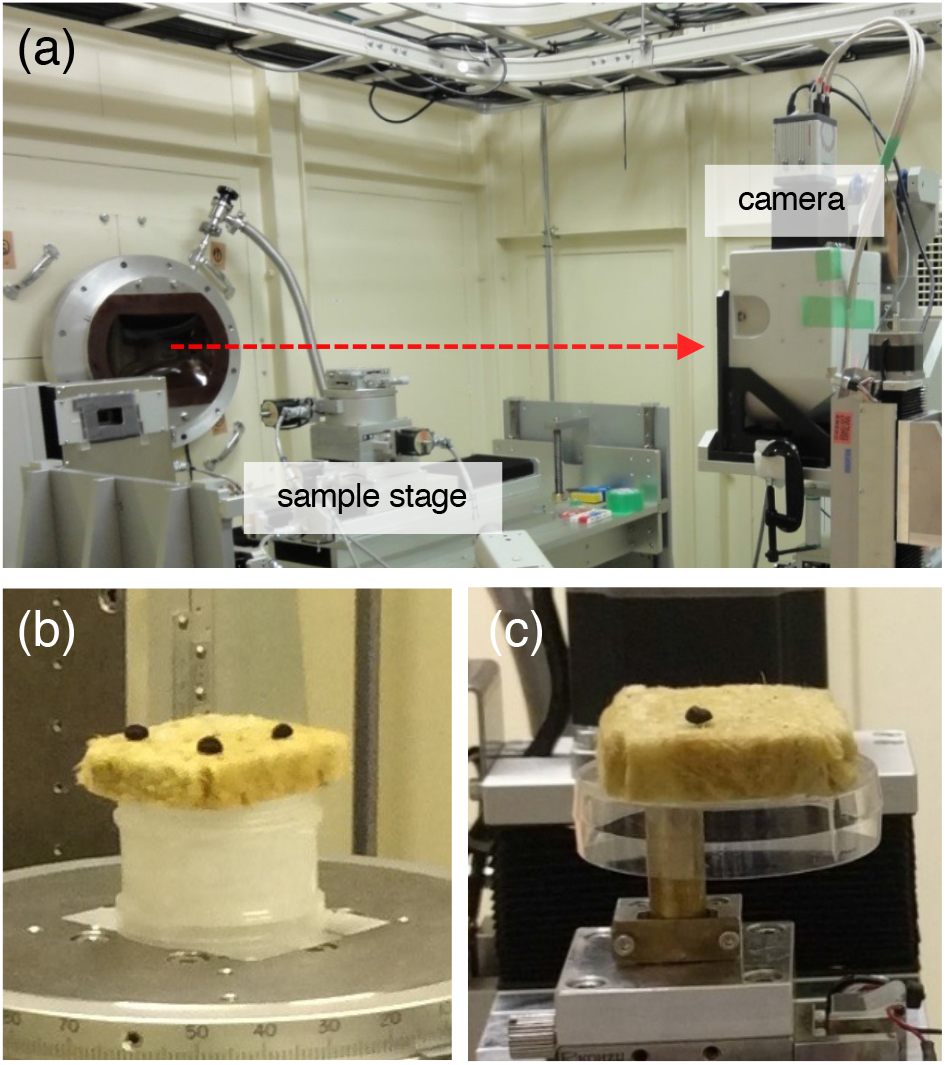
Experimental set up of the beamline BL20B2. A broken arrow (a) indicates the X-ray path in the Hutch 3. A rockwool slab placed on a rotation stage in the Hutch 3 (b) or the Hutch 1 (c). Seeds of morning glory were placed on the rockwool slabs as position markers.

**Fig. 3.**
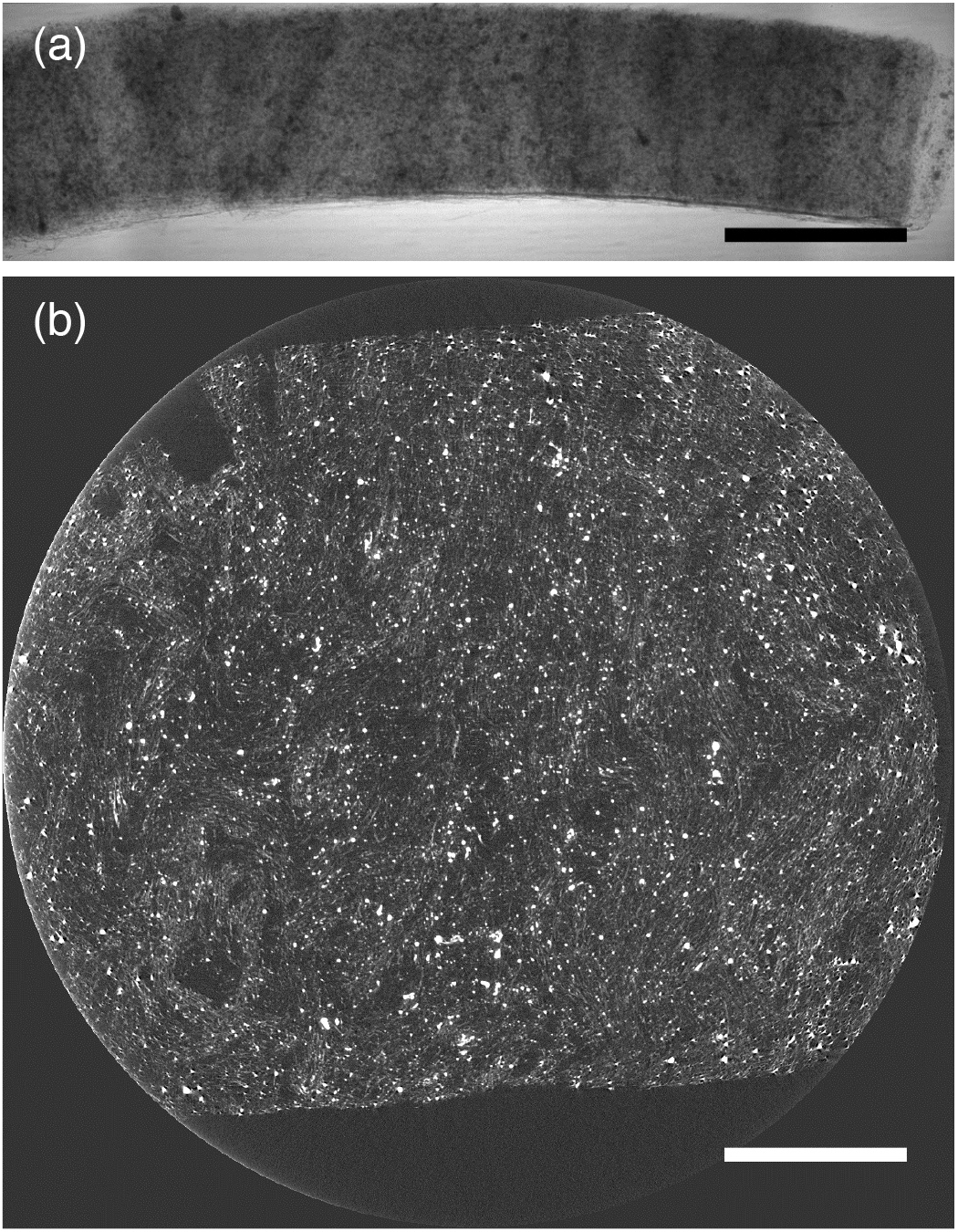
Two-dimensional projection of refraction contrast image of the rockwool slab (side view) (a) and a transverse tomographic slice of the rockwool slab (b). Scale bars = 1 cm.

### Optical microscopy

For microscopy of the cut surface of the root base and rockwool fibers, a multizoom microscope (AZ100M, Nikon, Tokyo, Japan) equipped with a CMOS camera DS-Fi3 (Nikon, Tokyo, Japan) was used.

## Results and discussion

### Identification of individual roots in a tomographic slice obtained by the observation at Hutch 3

After removing the shoot (upper) part of the plant, we identified positions of the plants under a multizoom microscope, finding cut surfaces of root or hypocotyl of the plants. Referring the position of the plant on the micrograph (Fig. 4a and b), we identified the root in a tomographic slice (Fig. 4c). Cross section of a root (or hypocotyl) often appeared ring-shape in a transverse tomographic slice because large cavity is often formed inside the root possibly when the plant was completely dried (Fig. 4c). The root system of Arabidopsis is a taproot system and is mainly composed of the primary root as the taproot and the secondary root as the lateral root. Shape of a primary root grown vertically downward in the rockwool slab was visualized in a longitudinal tomographic slice (Fig. 4d).

**Fig. 4.**
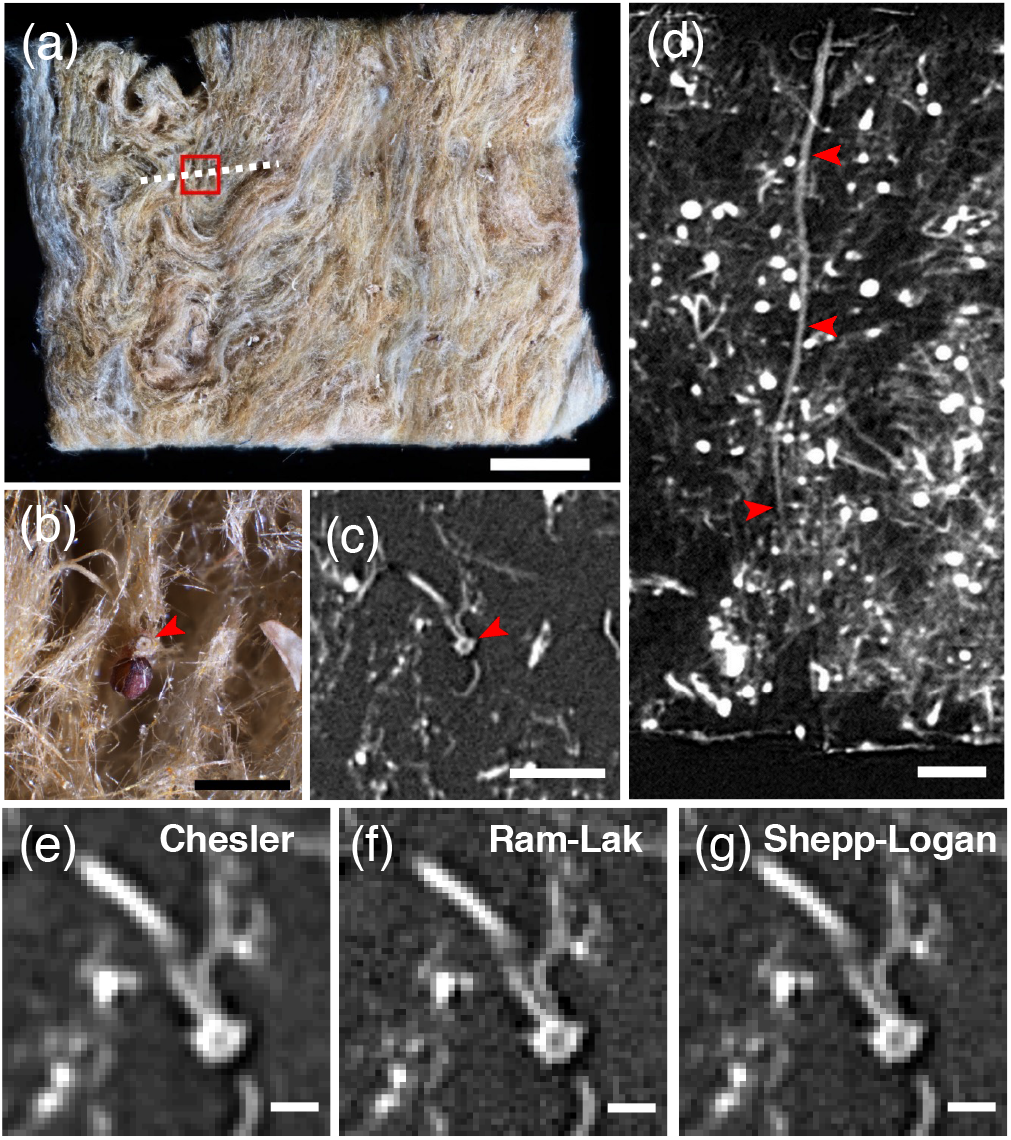
Identification of individual roots in a tomographic slice. (a) A multizoom micrograph of an entire rockwool slab. A red rectangle shows the area magnified in (b). A white dashed line shows the position from where the longitudinal tomographic slice shown in (d) was obtained. (b) A magnified microscopic view showing a cut surface of the base of the root found on the surface of the rockwool slab (red arrowhead). A dark-brown round-shape thing located nearby is a released seed coat, which frequently helps to find the position of the root. (c) A transverse tomographic slice showing the same position as shown in (b). White ring-shape object (red arrowhead) shows the cross section of the root. (d) A longitudinal tomographic slice obtained at the position indicated in (a) demonstrating longitudinal continuity of a primary root (red arrowheads). (e) – (g) Comparison of the tomographic slices reconstructed using different filters. (e) Chesler. (f) Ram-Lak. (g) Shepp-Logan. (c) and (d) are actually made using Chesler filter. Scale bars = 1 cm (a), 1 mm [(b) – (d)].

Next, we compared the appearance of the root in transverse tomographic slices when three different filters, Chesler, Ram-Lak, and Shepp-Logan filter were used during tomographic reconstruction. When reconstructed using Chesler filter in the case of data obtained at Hutch 3, background noise was reduced while root boundary became obscure (Fig. 4e). When reconstructed using Ram-Lak and Shepp-Logan filter, root boundary appeared clearer compared with Chesler. However, because Ram-Lak filter gives root boundary smoother than Shepp-Logan filter (Fig. 4g) when carefully compared between them, we have finally chosen to use Ram-Lak filter (Fig. 4f).

### Three-dimensional volume imaging of an individual root by the observation at Hutch 3

Shapes of the roots were roughly visualized by automatically drawing of isosurface (volume) models based on the voxel value (Fig. 5a and b). A considerable amount of fibrous and particulate fragmentary structures, which are due to component materials of the rockwool, were observed around the root besides the root itself (Fig. 5a and b). Some of these structures could be removed using ”Delete small pieces” function of the isosurface command of IMOD software (Fig. 5c).

**Fig. 5.**
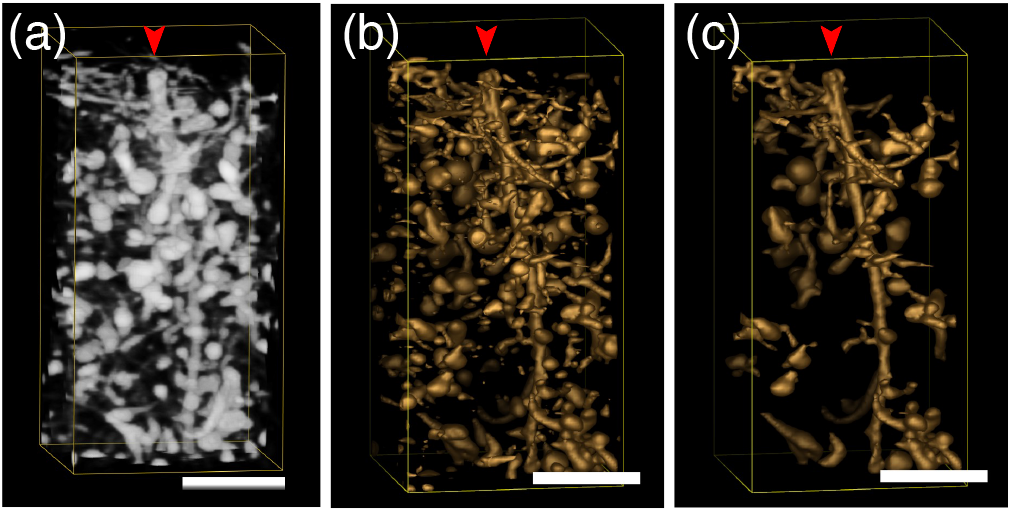
Three-dimensional volume imaging of individual root and determining its threshold. (a) A volume model made by using Volume Viewer of UCSF Chimera. (b) An isosurface model made by using isosurface command of IMOD software. (c) The same region where ”Delete small pieces” function of the isosurface command of IMOD software was applied. Scale bars = 1 mm.

Isosurface models change depending on the threshold level, above which all voxels are enclosed by the surface. For a quantitative approach, it is necessary to find out the appropriate threshold level. The actual cross-sectional area of the root at its cut surface was measured first (Fig. 6a). Then the cross-sectional areas of the isosurface models were measured at the same position as of Fig. 6a when the threshold level was changed from 70 to 76 (Fig. 6b-g). In this case, the closest value was achieved when the threshold level was 73. Therefore, we can conclude that the isosurface model closest to the actual root is the one demonstrated in Fig. 6f.

**Fig. 6.**
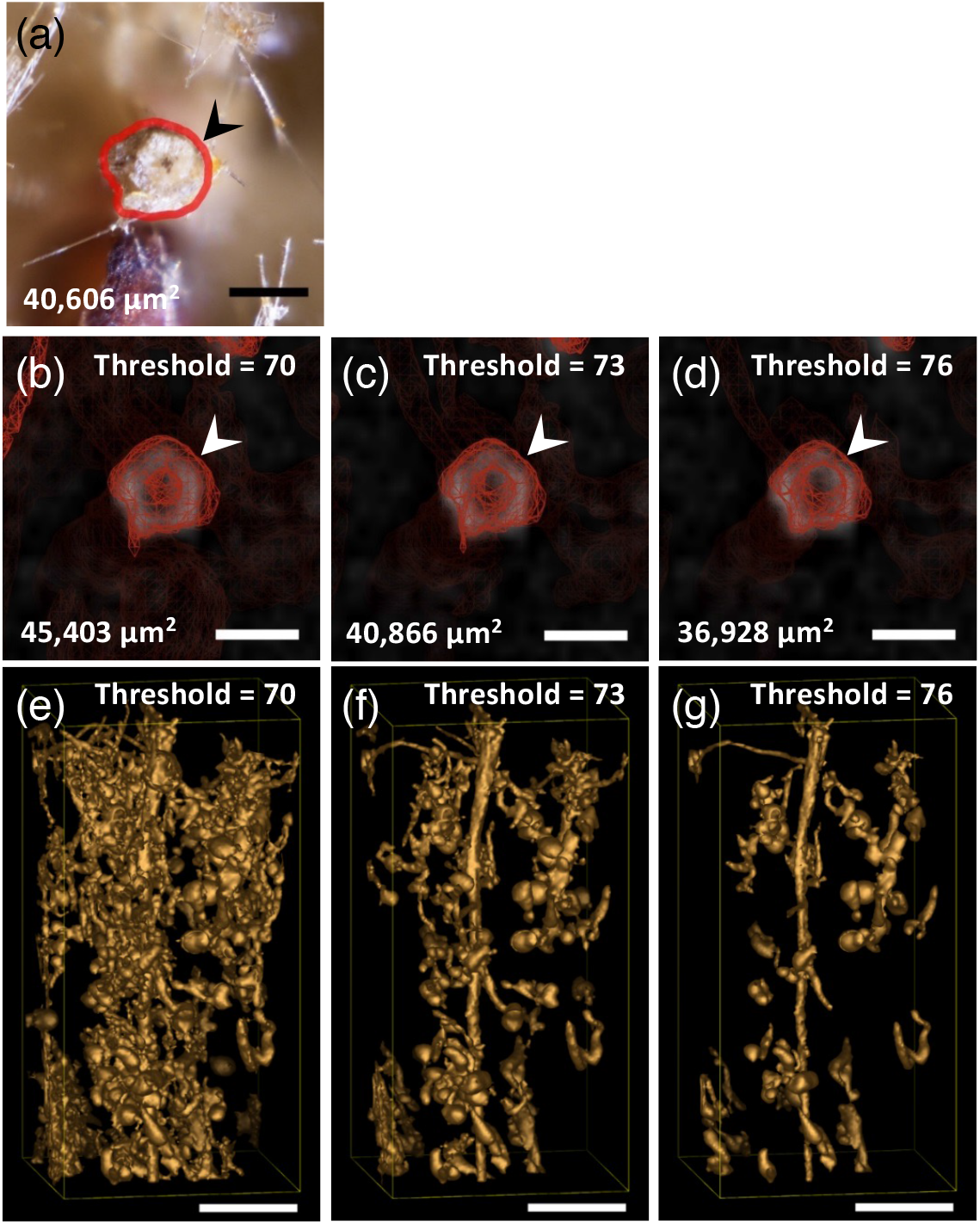
Finding the appropriate threshold level for isosurface rendering of the root. (a) A magnified multizoom micrograph showing the cut surface of the base of a root. A closed contour drawn in red shows the boundary of the root surface (arrowhead). The value is the actual cross-sectional area of the root surrounded by the contour. (b), (c), and (d) Cross sections of the isosurface models of the same root as shown in (a) (arrowheads) and their cross-sectional area values at the same position as shown in (a). (e), (f), and (g) Lateral views of the isosurface rendering which correspond to (b), (c), and (d), respectively. The threshold level was changed from 70 (b) and (e), 73 (c) and (f), to 76 (d) and (g). Scale bars = 200 μm [(a) – (d)], 1 mm [(e) – (g)].

However, even if the threshold level is fixed at the most appropriate value, unexpected connections between the root and surrounding rockwool materials were frequently observed. Also, unexpected disconnection of signals of the root occurred due to local attenuation of signals possibly at the thinner part of the root. These problems made it difficult to construct a 3D model of an entire root system only by making an isosurface model at one representative threshold value.

Instead, 3D wireframe models of roots were made, which is useful to measure the root length as well as to obtain other parameters representing morphology of roots, such as tortuosity, by manually tracing the signal of the root from the base toward the tip, simultaneously referring isosurface models adjusting around the appropriate threshold level (Fig. 7).

**Fig. 7.**
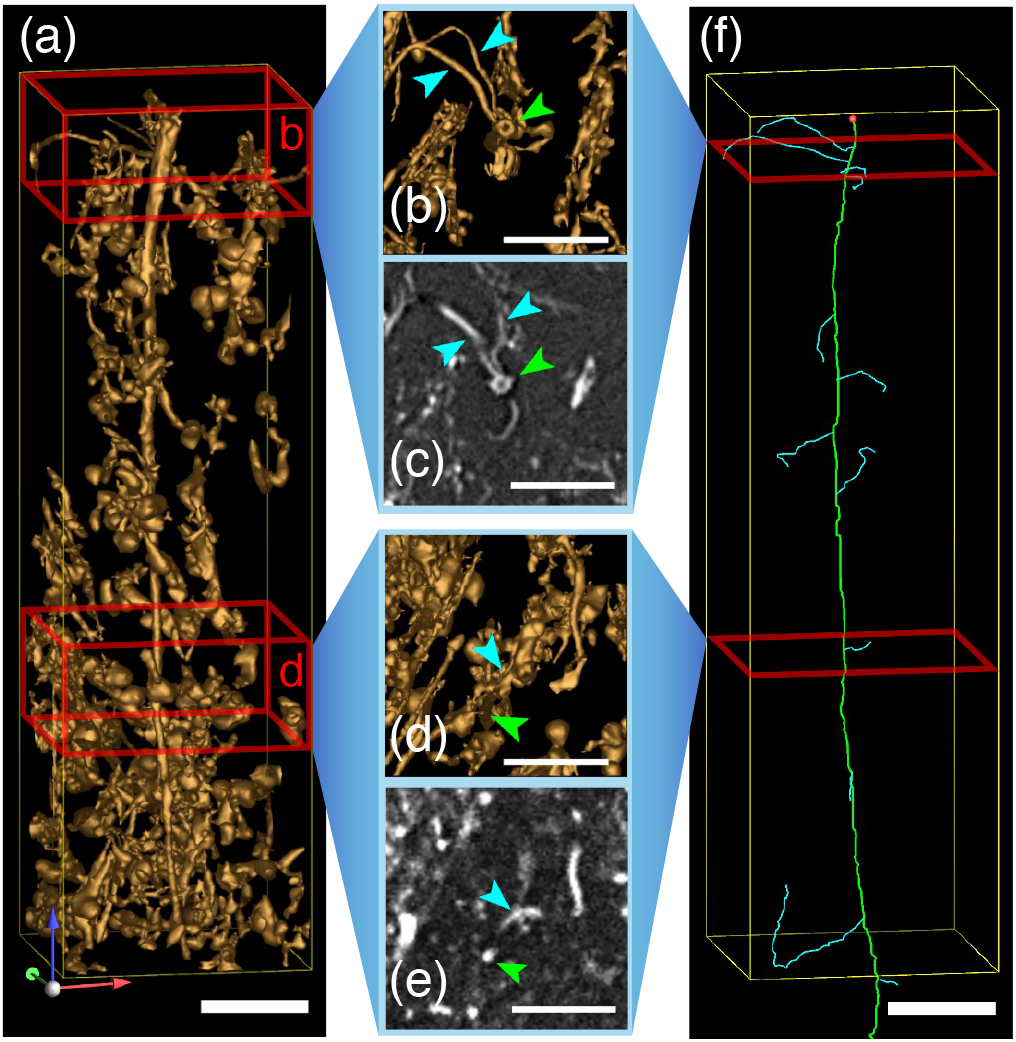
Making wireframe models of roots. (a) Lateral view of an isosurface model of the basal part of the root. Upper or lower red cuboid indicates the region shown in (b) or (d), respectively. (b) and (d) Transverse views of isosurface models from above corresponding to the regions indicated in (a). (c) and (e) Transverse tomographic slices at the positions corresponding to the regions indicated in (b) and (d), respectively. Green arrowheads show the primary root. Blue arrowheads show the secondary root. (f) A wireframe models of roots. Green contour shows the primary root. Blue contours show the secondary root. Scale bars = 1 mm.

Nevertheless, in many cases, unexpected disconnection of signals of the root frequently occurred, which made it difficult to trace entire part particularly of the secondary roots only by using Hutch 3 data. Therefore, we also tested Hutch 1 which enabled observation at higher spatial resolution.

### Observation of roots at higher spatial resolution at Hutch 1

In the case of data obtained at Hutch 1 where background noises were more noticeable (Fig. 8d and e), Chesler filter was used during tomographic reconstruction. Appearance of the transverse sections of the root is much clear in the tomographic slices of a reconstruction obtained by the observation at Hutch 1 (Fig. 8d and e) when compared with those obtained by the observation at Hutch 3 (Fig. 8a and b). Cavity formed inside the root can be clearly seen in the slice of the Hutch 1 reconstruction (Fig. 8d and e). An isosurface model of the root is much clearer when using the data obtained at Hutch 1 (Fig. 8f) than the data obtained at Hutch 3 (Fig. 8c). Rockwool fibers around the root were separated from the root and can be clearly seen by isosurface modeling using Hutch 1 reconstruction (Fig. 8f) although they were not separated from the root and coalesced into an agglomerated structure by isosurface modeling using Hutch 3 reconstruction (Fig. 8c).

**Fig. 8.**
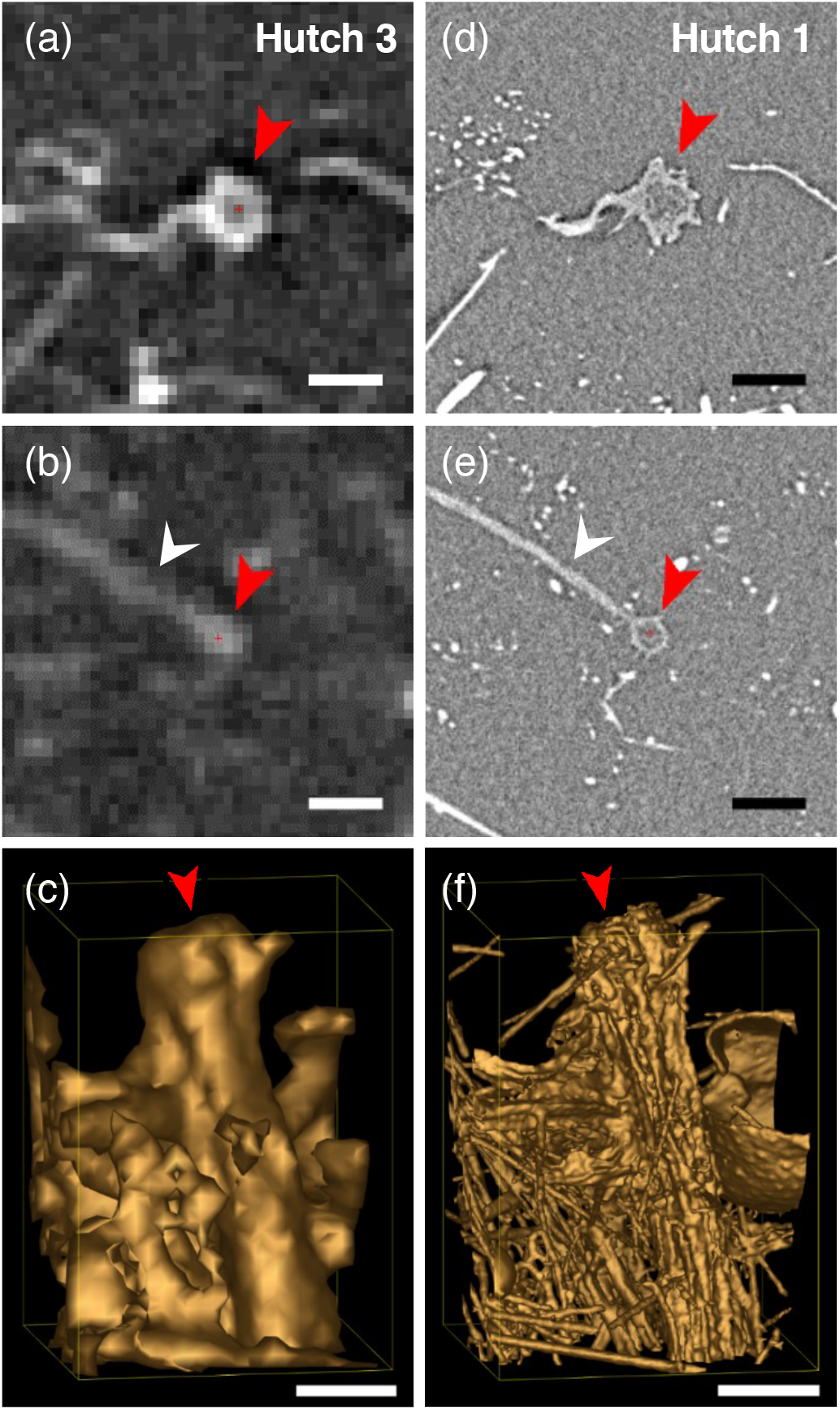
Comparison of the observations of the same root between Hutch 3 and Hutch 1. (a) and (b) Transverse tomographic slices obtained by the observation at Hutch 3. Effective pixel size = 25.5 μm pixel^-1^. (d) and (e) Transverse tomographic slices obtained by the observation at Hutch 1. Effective pixel size = 2.76 μm pixel^-1^. Red arrowheads show the transverse views of the root. (a) and (d) Transverse tomographic slices, which shows a thicker part of the primary root, obtained near the base of the root. (b) and (e) Transverse tomographic slices, which shows a thinner part of the primary root with a secondary root (white arrow heads), obtained at 4.6 mm below the base of the root. (c) and (f) Isosurface models using data obtained at Hutch 3 (c) or Hutch 1 (f). Red arrowheads show the base of the root. Scale bars = 200 μm.

Observation at Hutch 1 revealed detailed structures of the secondary root and component materials of the rockwool. The secondary root, which is generally thinner than the primary root, and rockwool fibers were distinguished by isosurface modeling using Hutch 1 data (Fig. 9b) while those structures were agglomerated in that of Hutch 3 (Fig. 9a). A fibrous form of rockwool observed in the Hutch 3 reconstruction does neither appear to be straight nor smooth but has an amorphous shape (Fig. 9a). On the other hand, a fiber observed in the Hutch 1 reconstruction appears to be straight and smooth (Fig. 9b and d) which is close to its shape observed under a multizoom microscope (Fig. 9e), indicating that the fibers recognized in the Hutch 1 data are considered to be individual fibers. Thickness of the isosurface model of the secondary root and a fibrous form of rockwool was measured and compared between in Hutch 3 and in Hutch 1 reconstructions (Table 1). As a result, thickness of a fibrous form of rockwool measured using the isosurface models of Hutch 3 data (78.0 ± 2.6 μm) was considerably thicker than that of Hutch 1 data (13.3 ± 0.1 μm), suggesting that a fibrous form of rockwool observed at Hutch 3 is considered to be a bundle of single rockwool fibers. And thickness of a fibrous form of a rockwool measured using the isosurface models of Hutch 3 data (78.0 ± 2.6 μm) was even thicker than the thickness of the secondary root measured using the isosurface models of Hutch 3 data (60.2 ± 7.6 μm), suggesting that this makes difficult to distinguish between the secondary root and a bundle of rockwool fibers in some cases. Thickness of a single rockwool fiber measured using the isosurface models of Hutch 1 data (13.3 ± 0.1 μm) was still thicker than its actual size measured using optical micrographs (9.1 ± 0.1 μm). Nevertheless, the isosurface model of a fibrous form of rockwool made using Hutch 1 data is considered to be a single fiber because its thickness (13.3 ± 0.1 μm) is still smaller than the twice of the thickness measured using optical micrographs (9.1 ± 0.1 μm). This result is comparable to, or even slightly higher than, the spatial resolution (voxel size, 28 μm) realized in the previous study done by Metzner et al. (2015) using Fraunhofer Development Center X-ray Technology (EZRT). As a result, the secondary roots could be successfully traced for the longer distance using Hutch 1 data than the Hutch 3 data due to higher spatial resolution (Fig. 10).

**Fig. 9.**
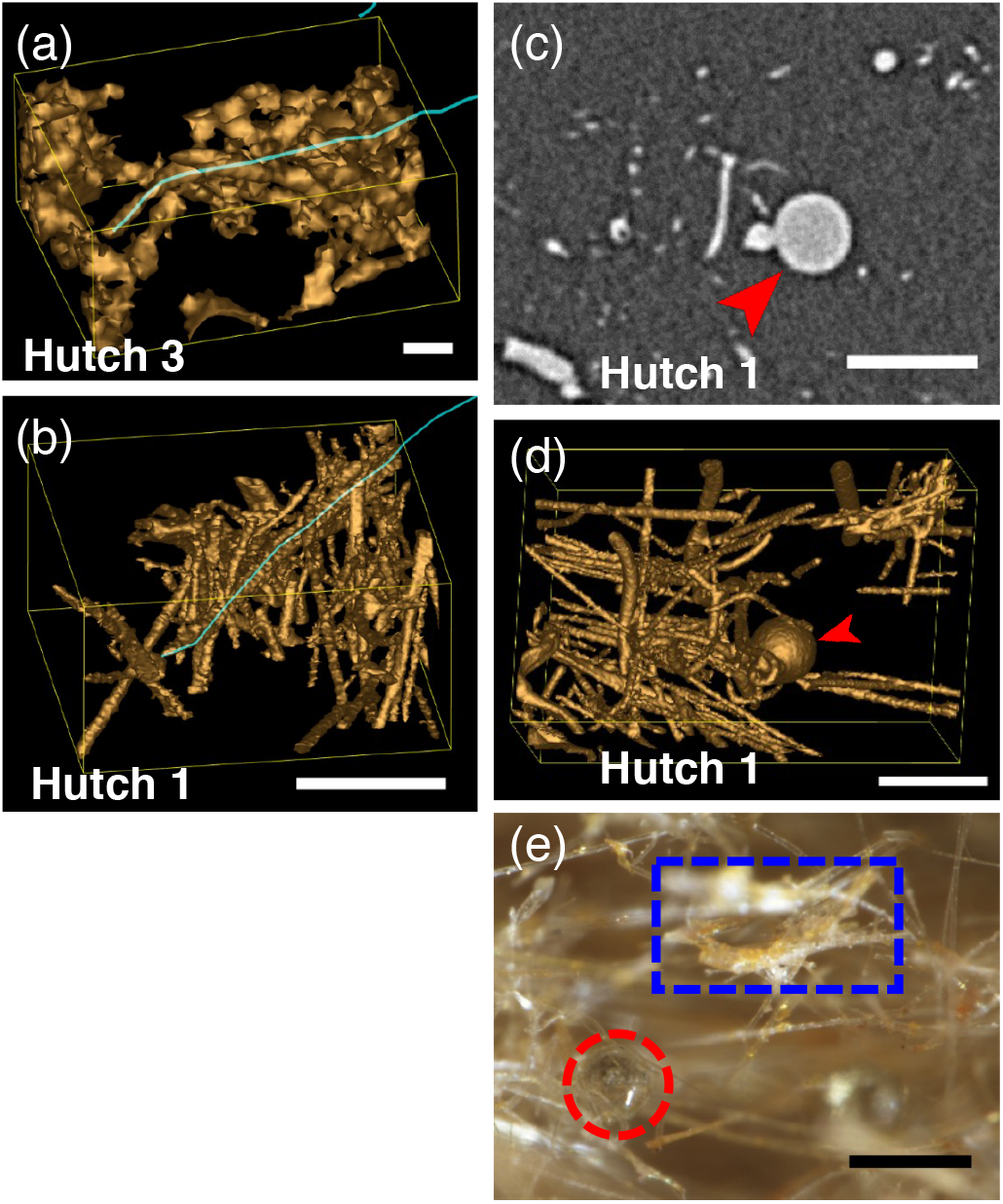
Detailed structures of the root and component materials of the rockwool revealed by the observation at Hutch 1. (a) and (b) Isosurface models of a fibrous form of rockwool and the secondary root obtained by the observation at Hutch 3 (effective pixel size is 25.5 μm pixel^-1^) (a) and Hutch 1 (effective pixel size is 2.76 μm pixel^-1^) (b), respectively. Blue open contours show wireframe models of the secondary roots. (c) and (d) A transverse tomographic slice (c) and its corresponding isosurface model (d) showing a globular shot structure of rockwool (arrowheads). (e) A multizoom micrograph showing an actual globular shot structure of rockwool (red broken circle) and ‘binder’ material agglutinating rockwool fibers (blue rectangular). Scale bars = 200 μm.

**Table 1.** Comparison of the thickness of the secondary root and a fibrous form of rockwool which was measured using the isosurface models of Hutch 3 and Hutch 1 as well as thickness of a rockwool fiber measured using multizoom micrographs

|  | Optical micrograph |  | Isosurface model |  |  |  |
| --- | --- | --- | --- | --- | --- | --- |
|  |  |  | Hutch 3 |  | Hutch 1 |  |
| | Mean ± SE ($\mu\text{m}$) | n | Mean ± SE ($\mu\text{m}$) | n | Mean ± SE ($\mu\text{m}$) | n |
| Fibrous form of Rockwool | 9.1 ± 0.1 | 35 | 78.0 ± 2.6 | 60 | 13.3 ± 0.1 | 60 |
| The secondary root | ND | | 60.2 ± 7.6 | 8 | 20.9 ± 1.4 | 8 |

ND, not determined.

**Fig. 10.**
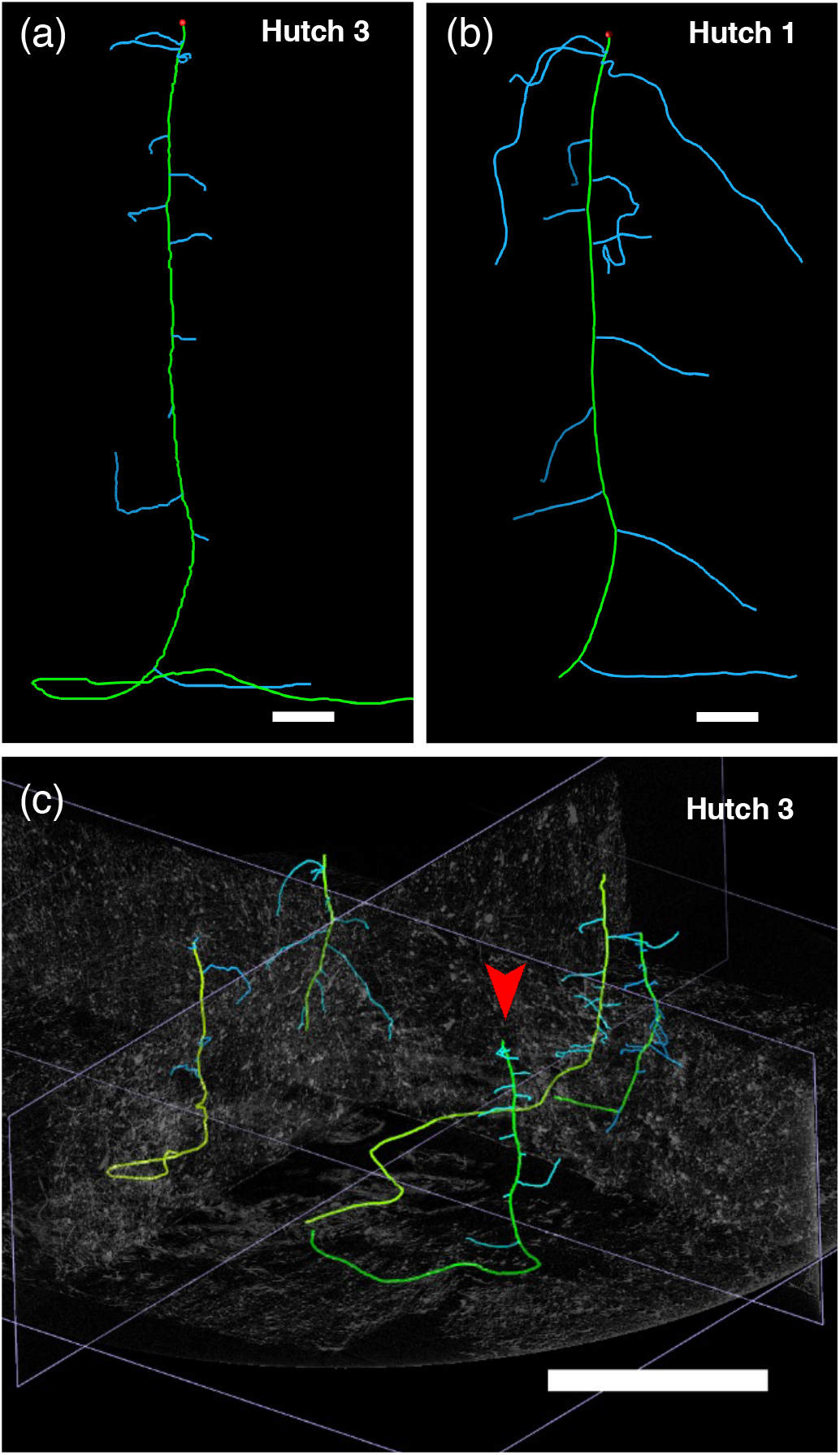
Comparison of wireframe models of the same root system drawn using Hutch 3 (a) and Hutch 1 (b) data. (c) Four root system models are seen in the wider view of the Hutch 3 data. Green contour shows the primary root. A red arrowhead shows the same root as in (a) and (b). In (b) apical part of the primary root missing because this missing part was out of the reconstructed volume data of the lower half. Blue contours show the secondary root. Scale bars = 1 mm (a) and (b), 10 mm (c).

In addition to the rockwool fibers, globular structures called ‘shot’, which was formed during the manufacturing of rockwool slabs [21], were clearly observed in a tomographic slice (Fig. 9c) as well as in isosurface models (Fig. 9d) of Hutch 1. These globular structures were also found in isosurface rendering (Fig. 4d) or in a tomographic slice (Fig. 5a) obtained using Hutch 3 data. Optical microscopy revealed a ‘shot’ structure as well as the material agglutinating rockwool fibers called ‘binder’, which is composed of resin and has amorphous structure (Fig. 9e). Existence of these component materials of rockwool slabs also made it difficult to distinguish the secondary root in the rockwool slab using Hutch 3 data particularly when the roots became thinner.

Figure 11 shows volume models of the root system made by using the volume viewer of UCSF Chimera adjusting the visible range of voxel value. In both cases of Hutch 3 and Hutch 1 data, structures related to the rockwool, such as globular ones, are visible in the broad configuration of visible range of voxel value (Fig. 11 a, c, and e) while morphology of the root system is more clearly visible in the narrow configuration (Fig. 11 b, d, and f). Globular structures have mostly disappeared particularly in the case of narrow configuration of Hutch 1 data (Fig. 11d and f). These results indicate that the materials of rockwool slabs have more opacity to X-ray, or, absorb more X-ray than the materials of the root.

**Fig. 11.**
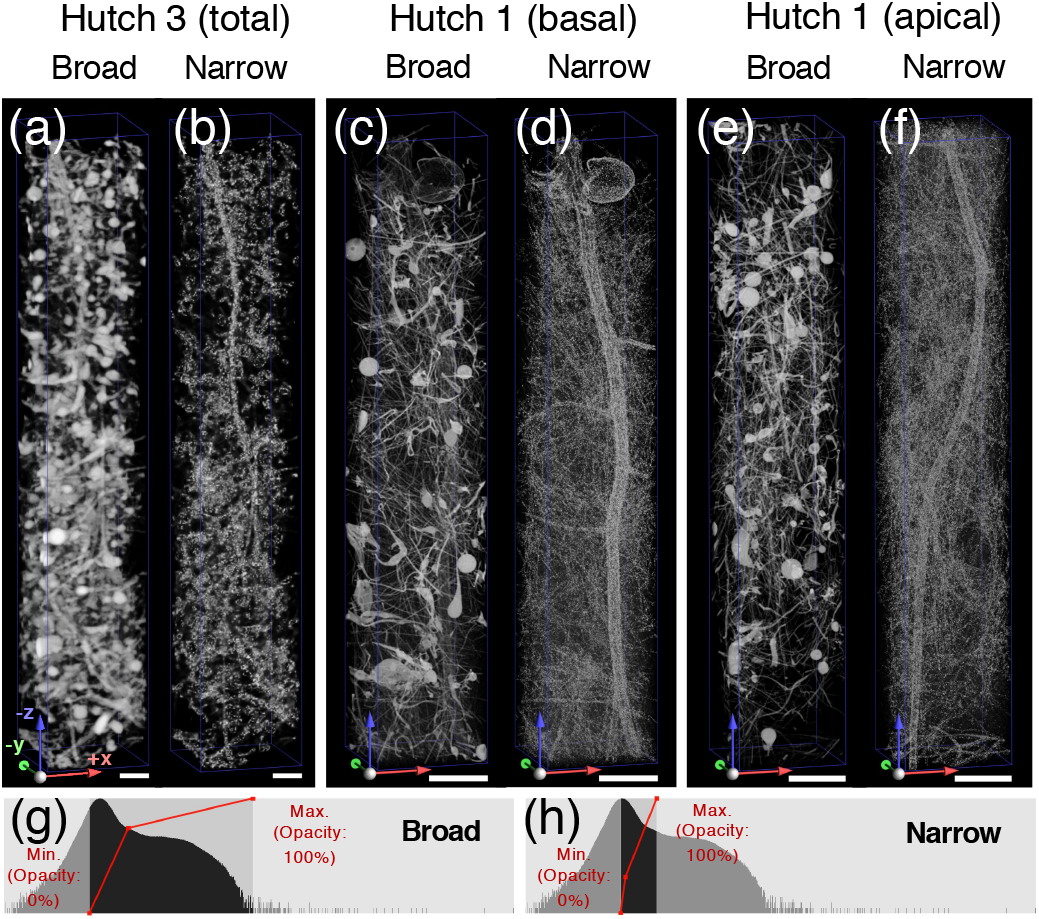
Visualization of the root system by using the volume viewer of UCSF Chimera adjusting the visible range of voxel value. (a) and (b) Volume models of the same root system using Hutch 3 data when visualized in the broad (a) or narrow (b) visible range of voxel value. (c) – (f) Volume models of the upper (c) and (d) or lower (e) and (f) part of the root using Hutch 1 data when visualized in the broad (c) and (e) or narrow (d) and (f) visible range of voxel value. Scale bars = 500 μm. (g) and (h) Examples of broad (g) or narrow (h) configuration of the visible range of voxel value which were actually applied to visualize the root volume model of (e) and (g) and (f) and (h), respectively. X axis shows voxel value. Y axis shows frequency. Visible areas are filled in black. Histograms show distribution of voxel values. Red line graphs show adjustment of opacity level.

The limit for the smallest observable root is a function of the quality of the image (signal-to-noise ratio) and resolution [6]. The smallest observable object is generally considered to be of a size of twice the spatial resolution and this may even not be sufficient if the image is noisy and background is not homogeneous [22]. In other words, the smallest observable object should be larger than 2 pixels / voxels. Because effective pixel size of Hutch 3 data is 25.5 μm pixel^-1^ and that of Hutch 1 data is 2.76 μm pixel^-1^, respectively, expected size of the smallest observable object is likely to be 51 μm pixel^-1^ and 5.52 μm pixel^-1^, respectively. Average size of the isosurface model of the secondary root made using Hutch 3 data was 60.2 μm (Table 1), which is comparable to the expected size of the smallest observable object at Hutch 3 (51 μm pixel^-1^). Even at this level, the spatial resolution, which is realized in the present system, is higher than the previous studies mentioned in the Introduction using an industrial CT scanner or a scanning system using microfocus X-ray source.

We can conclude that the spatial resolution of the observation at Hutch 1 is high enough to distinguish each rockwool fiber. Considering the difference between the actual thickness of the rockwool fiber (9.1 μm) and that of its isosurface model (13.3 μm), the actual thickness of the secondary root should be smaller than that of its isosurface model (20.9 μm). Further, it should be noted that dried roots analyzed in the present study had become thinner than those before drying.

A wireframe model of the root system drawn using Hutch 1 data demonstrated more extended architecture of the secondary roots as well as the primary root (Fig. 10b). On the other hand, that of Hutch 3 data demonstrated only the basal parts of the secondary roots while the architecture of the primary root was visualized entirely (Fig. 10a). Nevertheless, Hutch 3 data are useful because these provide an overview map for roughly comparing the growth of roots as well as for selecting individual roots for further detailed analyses of actual specimens of the Space Seed experiment. Together, precise determination of the minimum thickness of the secondary root observable remains to be done for that study.

## Conclusions

We can conclude that the spatial resolution of the observation at Hutch 1 is high enough to distinguish individual rockwool fibers as well as the secondary roots having the thickness of its isosurface model at 20.9 μm on the average. This enabled us to distinguish between them and to trace the architecture of the Arabidopsis root system including the secondary roots as well as the primary root. In addition, the observation at Hutch 3 provides an overview map of the root system grown in the rockwool slab for the purpose of screening of specimens for further detailed analyses of individual root systems.

## Acknowledgements

We thank Ms. Chiaki Zenko for her technical assistance. The synchrotron radiation experiments were performed at the BL20B2 of SPring-8, with the approval of the JASRI (Proposal Nos. 2014B1225, 2016A1390, 2016B1371, 2017B1225, 2018B1182, 2019A1130, 2019B1339 and 2020A1264).

## Funding

This work was partly supported by MEXT KAKENHI [grant number 15K11914 to I. K.] and 2020 Front loading research grant funded by Japan Aerospace Exploration Agency (JAXA), Institute of Space and Astronautical Science (ISAS) Expert Committee for Space Environment Utilization Science.

## Notes

### Competing Interest Statement

The authors have declared no competing interest.

## References

1. O’toole J C and Bland W L (1987) Genotypic variation in crop plant root systems. Adv. Agron. 41: 91–145.

2. Tran T T, Kano-Nakata M, Suralta R R, Menge D, Mitsuya S, Inukai Y and Yamauchi A (2015) Root plasticity and its functional roles were triggered by water deficit but not by the resulting changes in the forms of soil N in rice. Plant Soil 386: 65–76.

3. Kano M, Inukai Y, Kitano H and Yamauchi A (2011) Root plasticity as the key root trait for adaptation to various intensities of drought stress in rice. Plant Soil 342: 117–128.

4. Kano-Nakata M, Inukai Y, Wade L J, Siopongco J D L C and Yamauchi A (2011) Root development, water uptake, and shoot dry matter production under water deficit conditions in two CSSLs of rice: Functional roles of root plasticity. Plant Prod. Sci. 14: 307–317.

5. Faget M, Nagel K A, Walter A, Herrera J M, Jahnke S, Schurr U and Temperton V M (2013) Root-root interactions: extending our perspective to be more inclusive of the range of theories in ecology and agriculture using in-vivo analyses. Ann. Bot. (Lond.) 112: 253–266.

6. Flavel R J, Guppy C N, Tighe M, Watt M, Mcneill A and Young I M (2012) Non-destructive quantification of cereal roots in soil using high-resolution X-ray tomography. J. Exp. Bot. 63: 2503–2511.

7. Rogers E D, Monaenkova D, Mijar M, Nori A, Goldman D I and Benfey P N (2016) X-ray computed tomography reveals the response of root system architecture to soil texture. Plant Physiol. 171: 2028–2040.

8. Mairhofer S, Zappala S, Tracy S, Sturrock C, Bennett M J, Mooney S J and Pridmore T P (2013) Recovering complete plant root system architectures from soil via X-ray μ-computed tomography. Plant Methods 9: 1–7.

9. Daly K R, Tracy S R, Crout N M J, Mairhofer S, Pridmore T P, Mooney S J and Roose T (2018) Quantification of root water uptake in soil using X-ray computed tomography and image-based modelling. Plant Cell Environ. 41: 121–133.

10. Gao W, Schlüter S, Blaser S R G A, Shen J and Vetterlein D (2019) A shape-based method for automatic and rapid segmentation of roots in soil from X-ray computed tomography images: Rootine. Plant Soil 441: 643–655.

11. Yoshida Y, Arita T, Otani J and Sawa S (2020) Visualization of Toyoura sand-grown plant roots by X-ray computer tomography. Plant Biotechnol. (Tokyo) 37: 481–484.

12. Teramoto S, Takayasu S, Kitomi Y, Arai-Sanoh Y, Tanabata T and Uga Y (2020) High-throughput three-dimensional visualization of root system architecture of rice using X-ray computed tomography. Plant Methods 16: 1–14.

13. Metzner R, Eggert A, Van Dusschoten D, Pflugfelder D, Gerth S, Schurr U, Uhlmann N and Jahnke S (2015) Direct comparison of MRI and X-ray CT technologies for 3D imaging of root systems in soil: potential and challenges for root trait quantification. Plant Methods 11: 1–11.

14. Karahara I, Suto T, Yamaguchi T, Yashiro U, Tamaoki D, Okamoto E, Yano S, Tanigaki F, Shimazu T, Kasahara H, Kasahara H, Yamada M, Hoson T, Soga K and Kamisaka S (2020) Vegetative and reproductive growth of Arabidopsis under microgravity conditions in space. J. Plant Res. 133: 571–585.

15. Karahara I, Yamauchi D, Uesugi K and Mineyuki Y (2015/4) Three-dimensional imaging of plant tissues using X-ray micro-computed tomography. Plant Morphol. 27: 21–26.

16. Yano S, Kasahara H, Masuda D, Tanigaki F, Shimazu T, Suzuki H, Karahara I, Soga K, Hoson T, Tayama I, Tsuchiya Y and Kamisaka S (2013) Improvements in and actual performance of the Plant Experiment Unit onboard Kibo, the Japanese experiment module on the international space station. Adv. Space Res. 51: 780–788.

17. Karahara I, Umemura K, Soga Y, Akai Y, Bando T, Ito Y, Tamaoki D, Uesugi K, Abe J, Yamauchi D and Mineyuki Y (2012) Demonstration of osmotically dependent promotion of aerenchyma formation at different levels in the primary roots of rice using a “sandwich” method and X-ray computed tomography. Ann. Bot. (Lond.) 110: 503–509.

18. Uesugi K, Hoshino M, Takeuchi A, Suzuki Y, Yagi N and Nakano T (2010) Development of fast (sub-minute) micro-tomography. AIP Conf. Proc. 1266: 47–50.

19. Kremer J R, Mastronarde D N and Mcintosh J R (1996) Computer visualization of three-dimensional image data using IMOD. J. Struct. Biol. 116: 71–76.

20. Karahara I, Suda J, Tahara H, Yokota E, Shimmen T, Misaki K, Yonemura S, Staehelin L A and Mineyuki Y (2009) The preprophase band is a localized center of clathrin-mediated endocytosis in late prophase cells of the onion cotyledon epidermis. Plant J. 57: 819–831.

21. Kitahara H (2015) Filamentation technology of rock wool. Nichias Technol. Rev. 368: 1–5.

22. Kaestner A, Schneebeli M and Graf F (2006) Visualizing three-dimensional root networks using computed tomography. Geoderma 136: 459–469.

